# DeepTRIAGE: Interpretable and Individualised Biomarker Scores using Attention Mechanism for the Classification of Breast Cancer Sub-types

**DOI:** 10.1101/533406

**Authors:** Adham Beykikhoshk, Thomas P. Quinn, Samuel C. Lee, Truyen Tran, Svetha Venkatesh

## Abstract

**Motivation:** Breast cancer is a collection of multiple tissue pathologies, each with a distinct molecular signature that correlates with patient prognosis and response to therapy. Accurately differentiating between breast cancer sub-types is an important part of clinical decision-making. Already, this problem has been addressed using machine learning methods that separate tissue samples into distinct groups, there remains unexplained heterogeneity within the established sub-types that cannot be resolved by the commonly used classification algorithms. In this paper, we propose a novel deep learning architecture, called `DeepTRIAGE` (Deep learning for the TRactable Individualised Analysis of Gene Expression), which not only classifies cancer sub-types with good accuracy, but simultaneously assigns each patient their own set of interpretable and individualised biomarker scores. These personalised scores describe how important each feature is in the classification of any patient, and can be analysed post-hoc to generate new hypotheses about latent heterogeneity.

**Results:** We apply the `DeepTRIAGE` framework to classify the gene expression signatures of luminal A and luminal B breast cancer sub-types, and illustrate its use for genes as well as the GO and KEGG gene sets. Using `DeepTRIAGE`, we calculate personalised biomarker scores that describe the most important features for classifying an individual patient as luminal A or luminal B. In doing so, `DeepTRIAGE` simultaneously reveals heterogeneity within the luminal A biomarker scores that significantly associate with tumour stage, placing all luminal samples along a continuum of severity.

**Availability and implementation:** The proposed model is implemented in Python using PyTorch framework. The analysis is done in Python and R. All methods and models are freely available from https://github.com/adham/BiomarkerAttend.

## 1 Introduction

Breast cancer is a collection of multiple tissue pathologies with a joint genetic and environmental aetiology, and is a leading cause of death among women worldwide. During the progression of cancer, inherited or acquired mutations in the DNA change the sequence (or amount) of the messenger RNA (mRNA) produced by the cell, thereby changing the structure (or amount) of functional protein. As such, mRNA can serve as a useful proxy for evaluating the functional state of a cell, with its abundance being easily measured by microarray or high-throughput RNA sequencing (RNA-Seq). Indeed, mRNA abundance has already been used as a biomarker for cancer diagnosis and classification (Golub *et al*. (1999); Bair and Tibshirani (2003)), cancer sub-type classification (Sørlie *et al*. (2001); Parker *et al*. (2009)), and for clustering gene expression signatures (Ben-Dor *et al*. (1999)). For a comprehensive comparison of the supervised and unsupervised methods used with gene expression data, see Pirooznia *et al*. (2008).

Despite advancements in the field, mRNA-based classifiers still present unique challenges. First, these data-sets contain many more features (10,000s) than samples (100s), a *p* ≫ *n* problem that is usually addressed by feature selection or feature engineering (Saeys *et al*. (2007); Kursa (2014)). Second, it is often difficult to interpret mRNA-based classifiers because the predictive genetic features do not necessarily make sense to biologist experts without explicit contextualisation. Third, the routine use of discriminative methods (e.g., support vector machines (Vanitha *et al*. (2015)) or random forests (Cai *et al*. (2015); Kursa (2014))) only provide information with regard to the importance of a feature for an entire class. For cancer data, this means that these methods cannot suggest the importance of a feature for a specific patient, but instead only provide such information at the level of cancer type or sub-type. This is important given that a substantial amount of heterogeneity remains unaddressed within cancer sub-types (Mayer *et al*. (2014); Parker *et al*. (2009)).

In this paper, we propose a deep learning method for the stratification of clinical samples that not only offers interpretability through feature importance at the level of the cancer sub-type, but also at the level of the individual patient. As such, our method offers a finer level of interpretation than existing methods by capturing the heterogeneity of samples within each sub-type. To achieve this goal, we use an *attention mechanism*, a deep learning technique first proposed for machine translation and automatic image captioning (Bahdanau *et al*. (2014); Xu *et al*. (2015)). Attention allows salient features to come dynamically to the forefront for each patient as needed. As a result, the global knowledge of the model, obtained from the discriminating classes, is enhanced by the local knowledge that each patient provides. In other words, the attention mechanism offers an insight into the model’s decision-making process by revealing a set of individualised importance scores that describe how important each feature is for the classification of that patient. Further analysis of these importance scores reveals valuable insights into sub-type heterogeneity that are not directly apparent in the unattended data.

Machine learning techniques have been successfully applied to gene expression data for decades. More recently, deep learning, especially unsupervised deep learning, has influenced several approaches to gene expression analysis. These unsupervised models have been used to learn meaningful abstractions of biology from unlabelled gene expression data. For example, Tan *et al*. (2016) extracted biological insights from the *Pseudomonas aeruginosa* gene expression compendium using shallow auto-encoders. Similarly, stacked auto-encoders have been adopted to capture a hierarchical latent space from yeast gene expression data (Chen *et al*. (2016)), showing that the first layer correctly captures yeast transcription factors, while deeper layers conform to the existing knowledge of biological processes. More related to our work, it has been shown that shallow denoising auto-encoders are capable of extracting clinical information and molecular signatures from the gene expression data of patients with breast cancer (Tan *et al*. (2015)). Later, Danaee *et al*. (2016) coupled representations obtained from stacked denoising auto-encoders with a traditional classifier to achieve discriminatory power, applying it to classify cancerous samples from healthy ones.

Although the classification of cancer is an important task, breast cancer is not a monolithic entity. It is comprised of distinct molecular sub-types, each with a distinct molecular signature that correlate with patient prognosis and response to therapy (Dai *et al*. (2015)). This is especially true when considering the luminal sub-types: luminal A cancers have a much better prognosis than luminal B cancers and can be treated with endocrine therapy alone (Dai *et al*. (2015)). Yet, luminal A remains one of the most diverse cancer sub-types in terms of its molecular signature and severity (Netanely *et al*. (2016)). As a case study, we focus our analysis on the difficult problem of classifying breast cancer sub-types based on gene expression signatures, and approach it using an end-to-end supervised deep learning model. Using publicly available data from The Cancer Genome Atlas (**TCGA**), we develop and apply a novel deep learning architecture, called `DeepTRIAGE` (Deep learning for the TRactable Individualised Analysis of Gene Expression). The `DeepTRIAGE` architecture achieves two key outcomes.

First, our architecture extends the attention mechanism to model data where the number of features is much larger than the number of observations.

Second, our architecture facilitates a new interpretation of feature importance by providing individualised patient-level importance scores. These patient-level importance scores can be analysed directly using multivariate methods to reveal *and describe* latent intra-class heterogeneity.

Taken together, our work establishes a computational framework for calculating interpretable and individualised biomarker scores that can accurately classify luminal sub-types, while simultaneously revealing *and describing* intra-class heterogeneity. Using `DeepTRIAGE`, we calculate personalised biomarker scores that describe the most important features for classifying an individual patient as luminal A or luminal B. In doing so, `DeepTRIAGE` simultaneously reveals heterogeneity within the luminal A biomarker scores that significantly associate with tumour stage, placing all luminal samples along a continuum of severity.

## 2 Materials and methods

### 2.1 Data acquisition

We retrieved the unnormalised gene-level RNA-Seq data for the TCGA breast cancer (BRCA) cohort (Weinstein *et al*. (2013)) using the `TCGAbiolinks` package (Colaprico *et al*. (2016)). After filtering any genes with zero counts across all samples, we performed an effective library size normalisation of the count data using `DESeq2` (Anders and Huber (2010)). To retrieve luminal A (**LumA**) and luminal B (**LumB**) sub-type status for the TCGA BRCA samples, we downloaded the “PAM50” labels from the supplementary data of Netanely *et al*. (2016). With 1148 PAM50 labels retrieved, we excluded patients that had more than one tumour sample sequenced. This left us with 528 LumA and 201 LumB samples. Of these, 176 LumA and 67 LumB samples were set aside as a test set.

### 2.2 Engineering annotation-level expression from genes

To reduce the dimensionality of the raw feature space, we transformed raw features from “gene expression space” into an “annotation space”. For this, we elected to use the Gene Ontology (**GO**) Biological Process and Kyoto Encyclopedia of Genes and Genomes (**KEGG**) annotation databases. Pathways with less than ten associated genes were removed before estimating pathway-level expression by taking the sum of counts for all genes in each pathway, as described and validated in Quinn *et al*. (2018). This results in 3942 **GO** features and 302 **KEGG** features that are then used for model training.

### 2.3 Model architecture

Gene expression data-sets usually have many more features than samples. Using a standard deep learning architecture with such data leads to a parameter explosion in the model that can cause over-fitting and reduce generalizability. Instead, our model aims to (i) reduce the number of free parameters of the model, (ii) encode global knowledge in the data by finding discriminatory features at cancer sub-type level (akin to a logistic regression), and (iii) encode local information provided by each individual patient using an attention mechanism. These innovations make the attention mechanism tractable for high-dimensional data.

#### 2.3.1 The DeepTRIAGE model

Let *d* be the dimension of the raw feature space and let **x**_*j*_ = [*x*_*j*1_, …, *x*_*jd*_] be the representation of sample *j* in this space. Our goal is to train a binary classifier that learns whether sample *j* belongs to the LumA class (*y*_*j*_ = 1) or the LumB class (*y*_*j*_ = 0).

Let **E**_*d×m*_ be an embedding matrix and let **e**_*i*_ = [*e*_*i*1_, …, *e*_*im*_] be the embedding vector for feature *i* ∈ {1: *d*}. Using Equation 1, we define 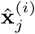, the *m*-dimensional embedded vector of feature *i* for sample *j*:

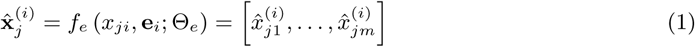

where *f*_*e*_, parametrised by Θ_*e*_, is the function that captures the relationship between the scalar value *x*_*ji*_ and the embedding vector **e**_*i*_. Note that the same embedding matrix is used for each sample.

Now, we can define a new representation for sample *j* using its embedded vectors, as shown in Equation 2:

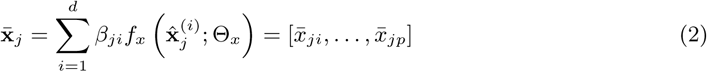

where 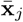 is a *p*-dimensional representation of the raw feature data, *f*_*x*_: ℝ^*m*^ →ℝ^*p*^ is parametrised by Θ_*x*_, and *β*_*ji*_ is a scalar value denoting the individualised **importance** that feature *i* has in the classification of sample *j*. Note that *m* ≪ *d* and usually *p* ≪ *m*. In other words, the dimensionality of the embedding space is much smaller than the dimensionality of the raw feature space, and the dimensionality of final representation 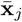 is smaller than the dimensionality of embedding space, allowing the attention mechanism to work successfully for such high-dimensional data. By viewing Equation 2 in a deep learning framework, one can interpret *β*_*ji*_ as the attention of sample *j* to feature *i* and compute it using Equations 3, 4, and 5:

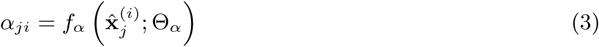

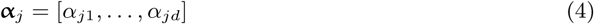

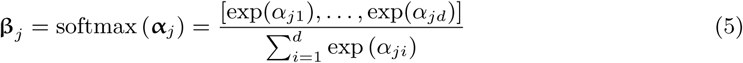

where *f*_*α*_: ℝ^*m*^→ℝ is parametrised by Θ_*α*_. Equation 5 is simply a normalisation to ensure that the attention weights sum to one for each sample.

Now, given the fixed size representation of sample *j* as 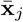, we can train the binary classifier *f*_*y*_ for LumA vs. LumB classification:

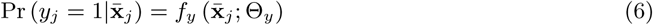

where Θ_*y*_ is the set of parameters for *f*_*y*_.

Given the embedding matrix **E**, we have defined our model through Equations 1-6. In the next section, we show how to learn the parameters of this model.

**Remark 1** There are different approaches to constructing the embedding matrix **E**. For instance: end-to-end learning with an unsupervised component added to the model, estimation using auto-encoders, or dimensionality reduction using PCA. We chose to use random vectors because it has been shown that their performance is comparable with the aforementioned techniques Bingham and Mannila (2001); Romero *et al*. (2016). Therefore, **e**_*i*_ is an *m*-dimensional random vector.

**Remark 2** There are many ways to compute the attention weights. We used a definition inspired by the concept of self-attention which means that the attention to a feature is only influenced by that feature (Vaswani *et al*. (2017)).

#### 2.3.2 Learning model parameters

In the previous section, we defined our model through Equation 1-6. Now we discuss how to specify its components {*f*_*e*_, *f*_*x*_, *f*_*α*_, *f*_*y*_} and how to learn their parameters {Θ_*e*_, Θ_*x*_, Θ_*α*_, Θ_*y*_}. Since we want to learn the model end-to-end, we choose these components to be differentiable.

In order to compute 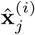, we capture the relationship between the feature value *x*_*ji*_ and the embedding vector **e***i* via multiplicative interaction using Equation 7. Therefore, Θ_*e*_ is a null set. One can, however, choose a more complex function.

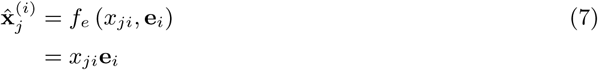

We choose *f*_*x*_ and *f*_*α*_ to be two feed-forward neural networks with weights Θ_*x*_ and Θ_*α*_ respectively. See Equations 8 and 9:

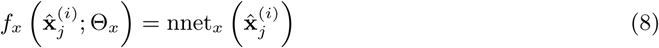

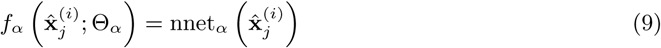

where both can be thought of non-linear transformation; nnet_*x*_: ℝ^*m*^→ℝ^*p*^ and nnet_*α*_: ℝ^*m*^→ℝ.

Given 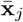, any differentiable classifier can be placed on top to predict the cancer sub-type (Equation 6). We use a feed-forward network with a sigmoid activation function in the last layer to calculate the probability of sample *j* belonging to a sub-type:

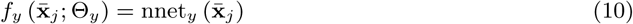

where Θ_*y*_ represents the weights of this network. To limit the model complexity, we choose *f*_*x*_ to be a single-layer neural network with tanh nonlinearity, *f*_*α*_ to be a network with one hidden layer and tanh nonlinearity, and *f*_*y*_ to be a network with one hidden layer, batch normalisation and tanh nonlinearity. Dropout with *p* = 0.5 is also applied to these three functions. Again, one can use more complex functions as long as they are differentiable.

Since all components are fully differentiable, the entire model can be learnt by minimising the log-loss function employing automatic differentiation and gradient-based methods. In this case, we used Adam optimiser (Kingma and Ba (2014)).

### 2.4 Analysis of importance scores

What we have described so far focuses on the discriminatory mechanism of our model. When viewed from the top, our proposed model is capable of separating cancer sub-types, like many other classification algorithms. However, one important distinction is that our model also *generates an individualised importance score for each feature at the sample-level*. This aspect is highly useful as it opens new opportunities for post-classification analyses of individual patients, making our method both hypothesis-testing and hypothesis-generating.

Given **β**_*j*_ = [*β*_*j*1_, *…, β*_*jd*_], where *β*_*ji*_ is the individualised importance score for sample *j* and feature *i*, we can construct an **importance score matrix B** by stacking **β**_*j*_ for all samples.

To detect emerging patterns within the individualised importance scores, we perform non-negative matrix factorisation (**NMF**) and principal component analysis (**PCA**) of the importance score matrix **B**. As a point of reference, we also perform an ordination of the raw feature space from 2.2. Note that all individualised per-sample importance scores were calculated on the withheld test set.

## 3 Results and Discussion

### 3.1 GINS1 drives luminal sub-type classification in test set

Table 1 shows the performance of the `DeepTRIAGE` model for luminal sub-type classification. When applying this model to Ensembl gene expression features, we obtain personalised biomarker scores that describe how important each gene is in predicting the cancer sub-type for each sample. The objective of `DeepTRIAGE` is to improve interpretability, not accuracy. Yet, this method appears to perform marginally better.

**Table 1:** This table shows the F1-score performance of the `DeepTRIAGE` attention model for luminal sub-type classification. We benchmark its performance as compared to a logistic regression and support vector machine (SVM), using both gene and gene set annotation features. From this, we see that our model, which adds a level of interpretability at the individual level, does not sacrifice classification accuracy. The objective of `DeepTRIAGE` is to improve interpretability, not accuracy. Yet, this method appears to perform marginally better.

|  | Logistic Regression | Linear SVM | DeepTRIAGE |
| --- | --- | --- | --- |
| **GO** (BP) annotations | 0.87 | 0.89 | 0.90 |
| **KEGG** annotations | 0.86 | 0.84 | 0.87 |
| Ensembl genes | 0.85 | 0.85 | 0.87 |

We can interpret the resultant importance score matrix directly using multivariate methods. Figure 1 shows the NMF factor which best discriminates between the breast cancer sub-types. Here, we see that a single gene, GINS1 (ENSG00000101003), contributes most to this factor. This gene has a role in the initiation of DNA replication, and has been associated with worse outcomes for both luminal A and luminal B sub-types (Nieto-Jiménez *et al*. (2017)). Interestingly, this is not a PAM50 gene, suggesting that our model is not merely re-discovering the PAM50 signature. We posit that the model performance, along with this biologically plausible result, validates its use for gene expression data.

**Figure 1:**
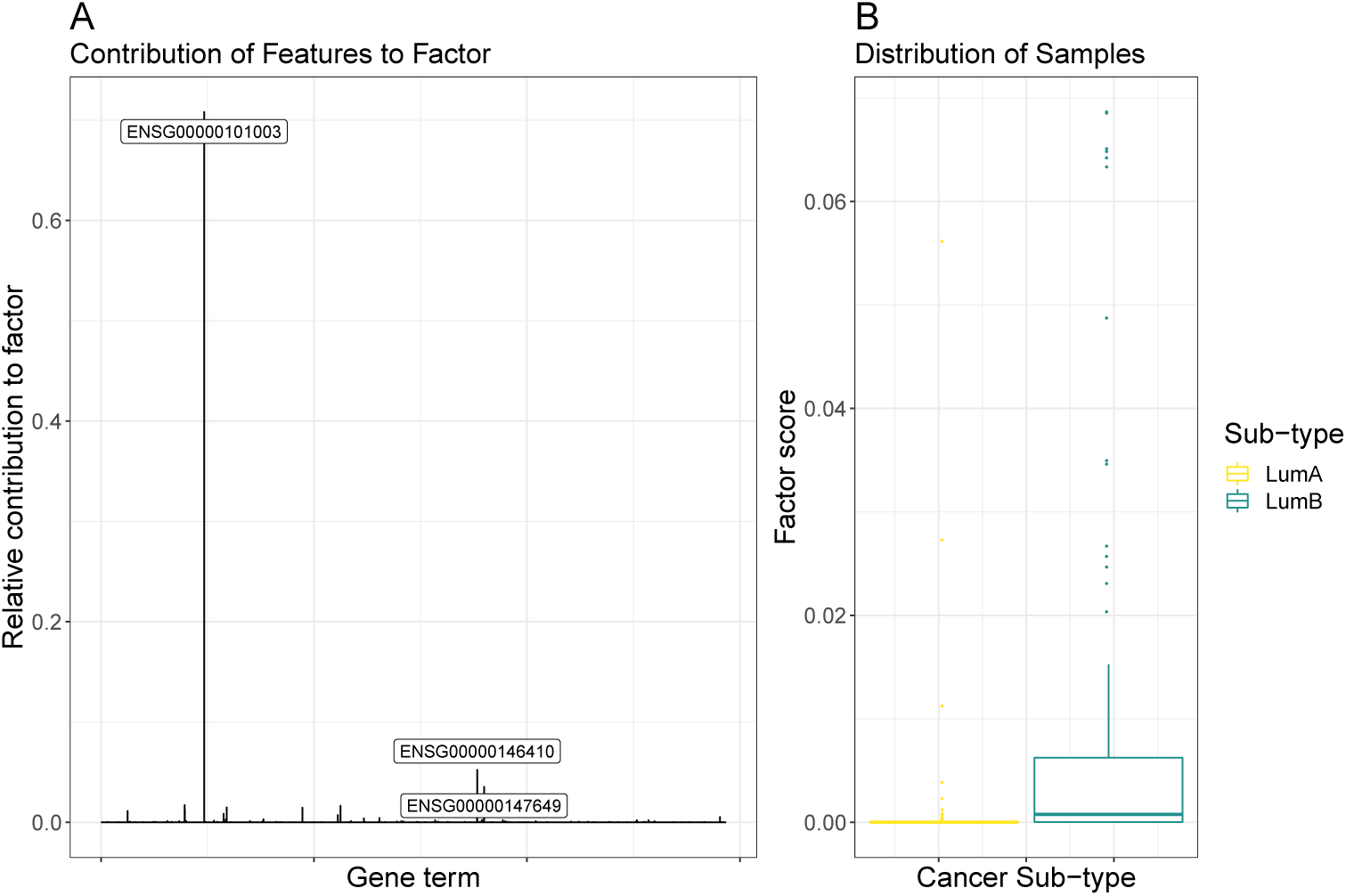
This figure presents the results of non-negative matrix factorisation applied to the importance score matrix computed from Ensemble gene expression data using DeepTRIAGE. Shown here is the factor which best discriminates between the two breast cancer sub-types. The left panel shows the relative contribution of each gene term to the most discriminative factor, with the top 3 components labelled explicitly. The right panel shows a box plot of the distribution of all samples across the composite factor score. This figure is produced using the test set only.

### 3.2 Kinetochore organisation associates with tumour severity within and between luminal sub-types

To reduce the number of features and to facilitate the interpretation of feature importance, we transformed the gene-level expression matrix into an annotation-level expression matrix using the Gene Ontology (**GO**) annotation set (cf. Section 2.2). Table 1 shows that **GO** annotation features perform as well as using gene features for all models. Although annotation features do not improve performance, they do improve the interpretability of the model by representing the data in a way that reflects domain-specific knowledge (Subramanian *et al*. (2005)). By applying `DeepTRIAGE` to the **GO** features, we obtain personalised biomarker scores that describe how important each **GO** term is in predicting the cancer sub-type for each sample.

Figure 2 shows the most discriminative NMF factor of the **GO**-based importance score matrix. The left panel shows the relative contribution of each term to this factor, while the right panel shows the distribution of samples with regard to this factor. From this, we see that a single factor cleanly delineates the luminal A samples from the luminal B samples, and is comprised mostly by the GO:0051383 (kinetochore organisation) gene set. Figure 3 shows a PCA of the same importance score matrix, along with a biplot of the 5 most variable **GO** terms, offering another perspective into the structure of the importance score matrix.

**Figure 2:**
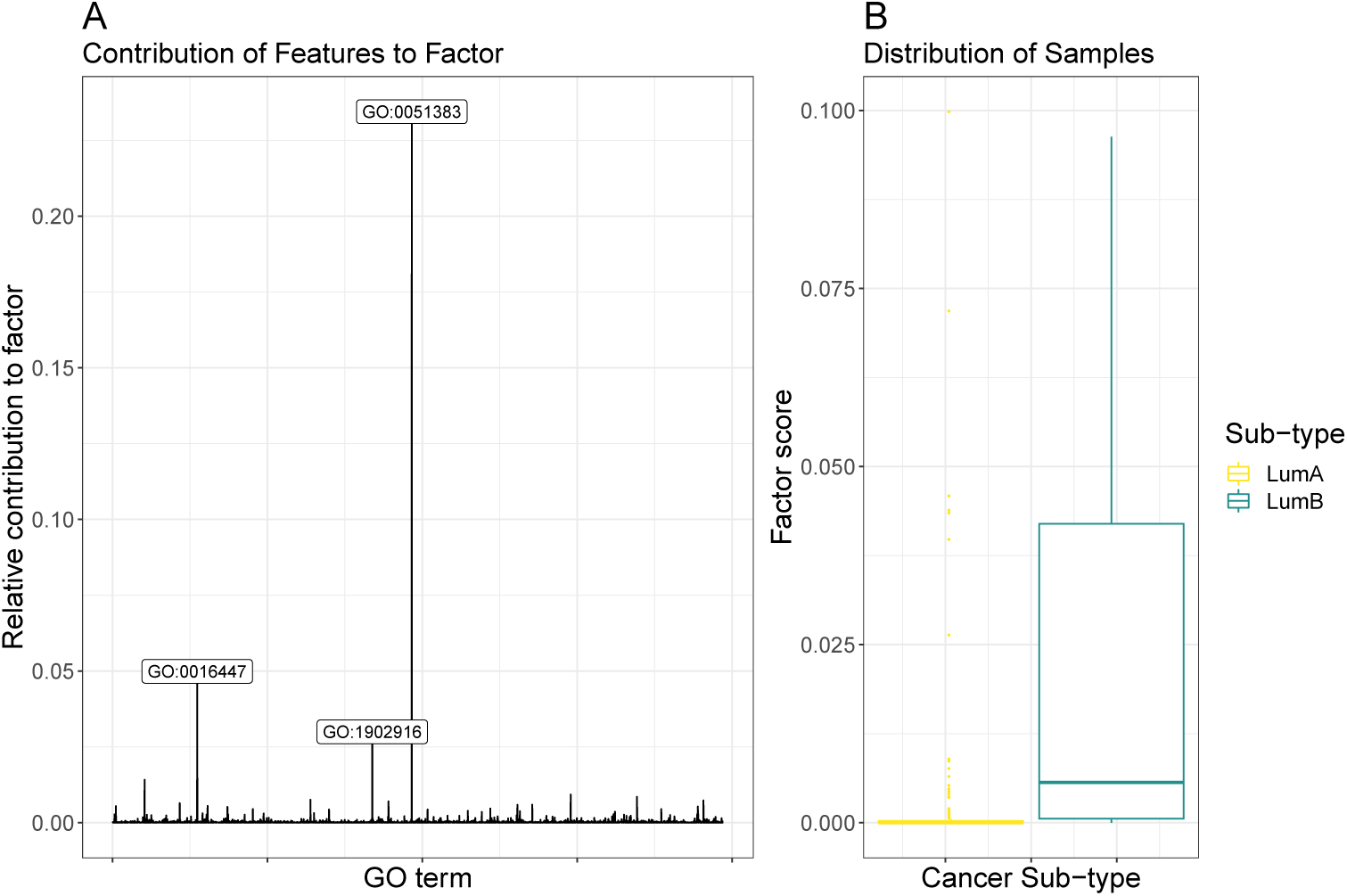
This figure presents the results of non-negative matrix factorisation applied to the GO-based importance score matrix. Shown here is the factor which best discriminates between the two breast cancer sub-types. The left panel shows the relative contribution of each **GO** term to the most discriminative factor, with the top 3 components labelled explicitly. The right panel shows a box plot of the distribution of all samples across the composite factor score. This figure is produced using the test set only.

**Figure 3:**
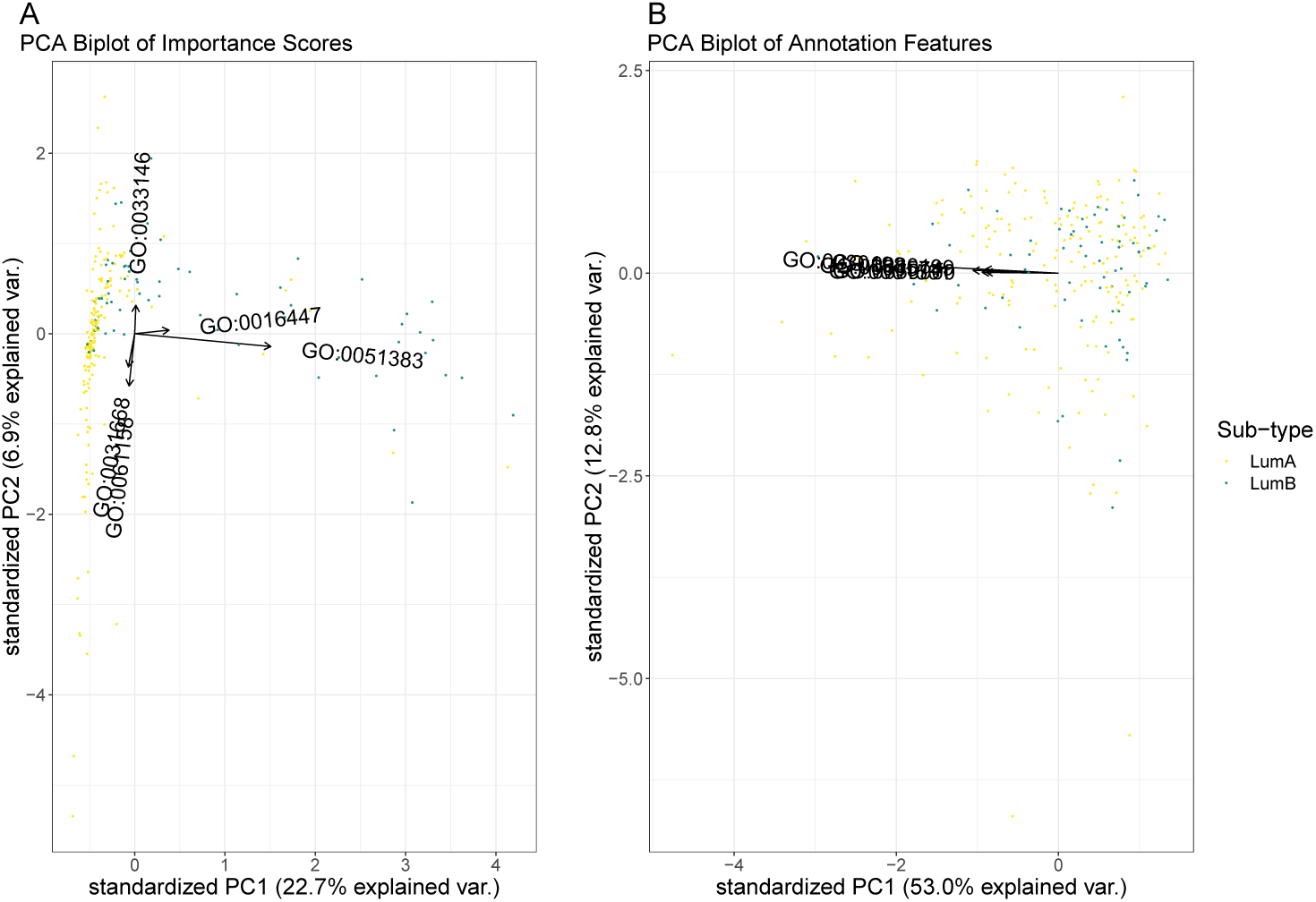
This figure shows a PCA biplot of the **GO**-based importance score matrix (left panel) and the **GO** annotation features (right panel), with the top 5 most variable terms labelled explicitly. For the importance scores, we see that the first principal axis describes much of the variance between the breast cancer sub-types, while the second principal axis describes much of the variance within the luminal A sub-type. By super-imposing the features as arrows, we can see which annotations best describe the origin of this variance. This level of structure is not evident when looking at the PCA biplot of the annotation feature space. This figure is produced using the test set only.

Both visualisations show that the kinetochore organisation gene set can meaningfully discriminate between the luminal A and luminal B cancer sub-types. This gene set contains 5 members: SMC4, NDC80, SMC2, CENPH, and CDT1. Figure 4 shows the expression of these genes in the test data, showing that the prioritised gene set contains genes with significant mean differences between the two sub-types (p-value < 0.01). Interestingly, only one of these (NDC80) is a member of the PAM50 gene set used to define the luminal A and B sub-types. The kinetochore organisation gene set is involved in the assembly and disassembly of the chromosome centromere, an attachment point for spindle microtubules during cell division. The dysregulation of this gene set would be expected to associate with luminal sub-typing because centromere instability drives genomic instability, and luminal B cancers are more unstable than luminal A cancers (as evidenced by Ki-67 staining (Inic *et al*. (2014)) and tumour severity). Indeed, NDC80 and CENPH dysregulation has already been associated with worse breast cancer outcomes, with luminal A exhibiting less centromere and kinetochore dysregulation in general (Zhang *et al*. (2016)).

**Figure 4:**
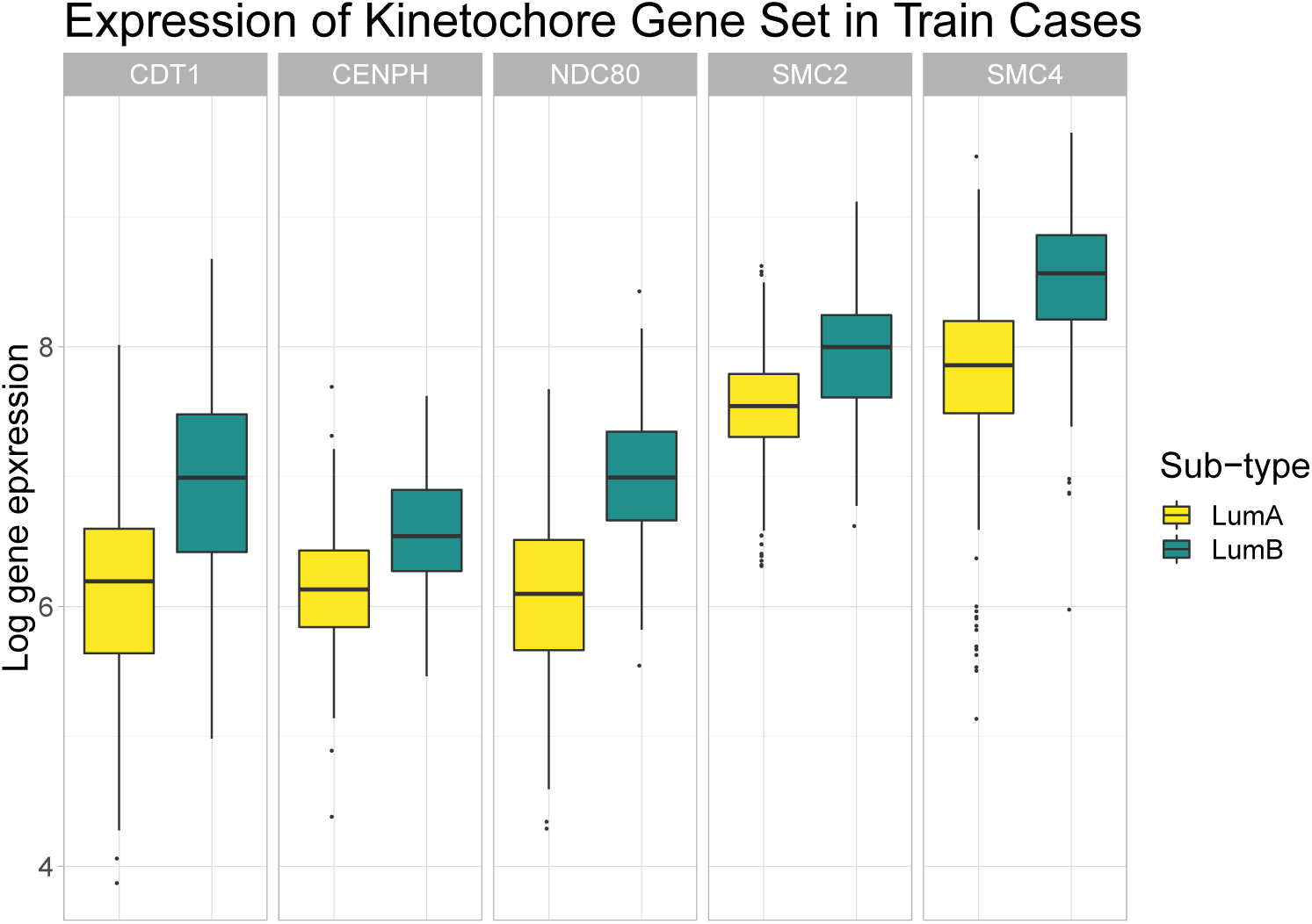
This figure shows the test set gene expression for 5 genes found within the GO:0051383 (kinetochore) gene set. Here, we see that all 5 genes are up-regulated in luminal B samples. This is relevant because our attention model prioritised this gene set when looking for feature importance within the breast cancer test set.

However, the real added value of our attention model is that it projects all samples according to a distribution of importance scores, implicitly revealing *and describing* heterogeneity within the cancer sub-types. While Figure 3 shows how GO:0051383 distinguishes between the luminal sub-types, it also shows how GO:0031668 (cellular response to extra-cellular stimulus) and GO:0061158 (3’-UTR-mediated mRNA destabilisation) explain much variance within the luminal A group. These axes are not arbitrary. A linear model predicting each PCA axis as a function of the tumour (T), node (N), and metastasis (M) stage (as nominal factors) *among the luminal A samples only*, reveals that small values in the first axis (PC1) significantly associate with the lower T stages, while large values significantly associate with the N2 stage (*p* < 0.05). Meanwhile, large values in the second axis (PC2) significantly associate with the T4 stage (*p* < 0.05). This suggests that the luminal A samples which are closest to luminal B samples in the PCA tend to be worse tumours. This is consistent with the literature which describes luminal B cancer as a more severe disease (Dai *et al*. (2015)), as well as Netanely et al’s observation that luminal cancers exist along a phenotypic continuum of severity (Netanely *et al*. (2016)). Thus, our method provides a biological explanation for some of the variance associated with the diagnostically-relevant differences in luminal sub-types. This level of resolution is not provided by the other machine learning algorithms used for RNA-Seq data, and is not evident in the ordination of the unattended **GO** annotation features (see Figure 3B).

### 3.3 DNA mismatch repair associates with tumour severity within and between luminal sub-types

We repeated the same analysis above using the Kyoto Encyclopedia of Genes and Genomes (**KEGG**) annotation set which organises genes according to canonical functional pathways (cf. Section 2.2). Like with **GO** annotations, the `DeepTRIAGE` model performed well with **KEGG** annotations (see Table 1). By applying `DeepTRIAGE` to the **KEGG** features, we obtain personalised biomarker scores that describe how important each **KEGG** term is for the classification of each patient.

The NMF and PCA ordination of the **KEGG**-based importance scores both show that hsa03430 (DNA mismatch repair) explains much of the inter-group variability (see Figures 5 and 6). This is expected to separate luminal A and B sub-types because errors in the DNA mismatch repair mechanism allow mutations to propagate, resulting in a more aggressive cancer. Yet, the PCA biplot shows that there exists a large amount of intra-class heterogeneity that is not explained by this pathway. Along this axis, we see a contribution by hsa04670 (Leukocyte transendothelial migration) and hsa04215 (Apoptosis), both relevant to tumour progression and metastasis. Again, these axes are not arbitrary. A linear model predicting each PCA axis as a function of the tumour (T), node (N), and metastasis (M) stage (as nominal factors) *among the luminal A samples only*, reveals that small values in both axes (PC1 and PC2) significantly associate with the T1 stage (*p* < 0.05). This suggests that the heterogeneity uncovered by the `DeepTRIAGE` architecture places patients along a diagnostically-relevant continuum of tumour severity. Again, this level of resolution is not provided by other machine learning algorithms and is not evident in the ordination of the unattended annotation-level data (see Figure 6B).

**Figure 5:**
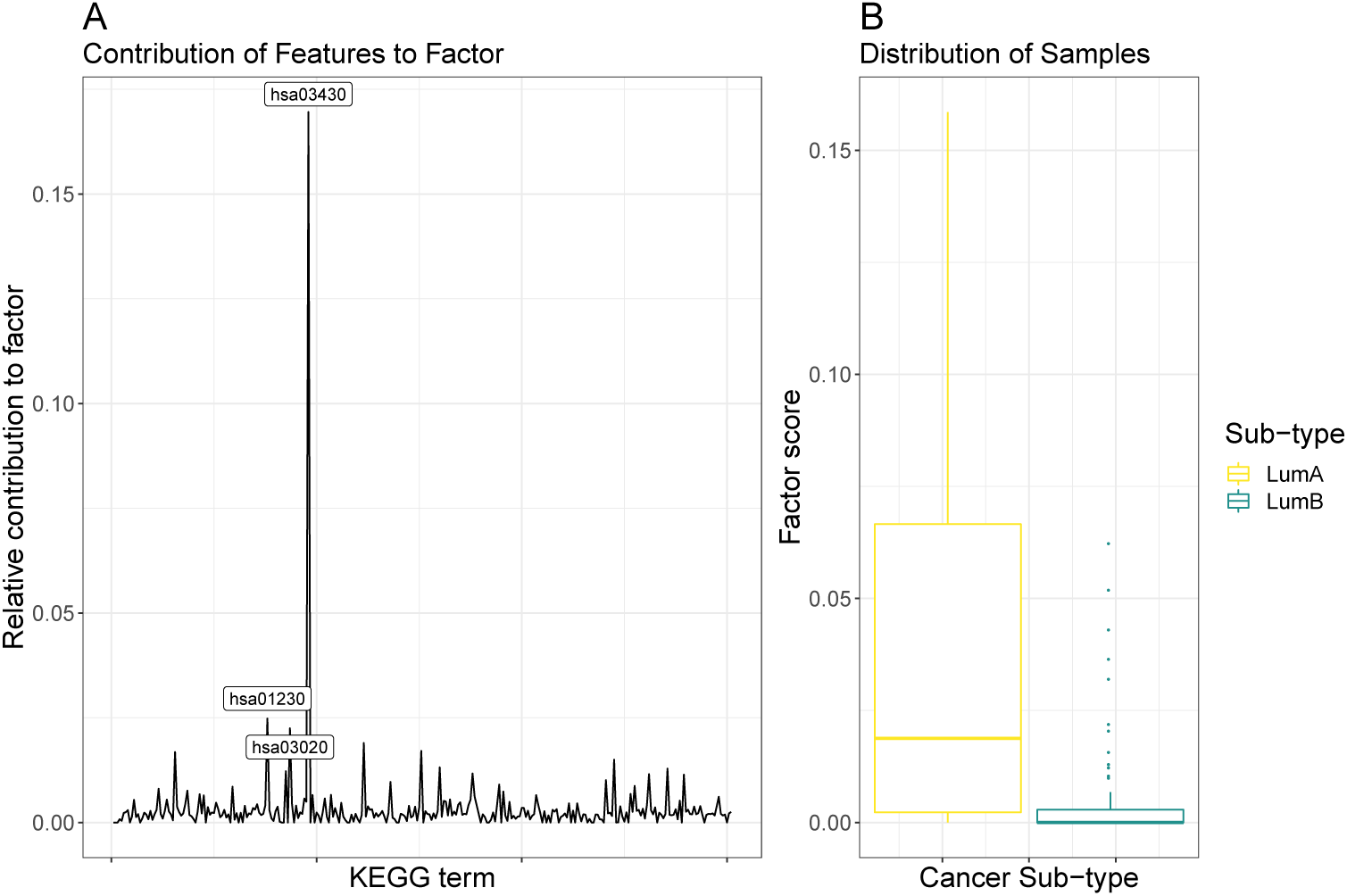
This figure presents the results of a non-negative matrix factorisation applied to the KEGG-based importance score matrix. Shown here is the factor which best discriminates between the two breast cancer sub-types. The left panel shows the relative contribution of each KEGG term to the most discriminative factor, with the top 3 components labelled explicitly. The right panel shows a box plot of the distribution of all samples across the composite factor score. This figure is produced using the test set only.

**Figure 6:**
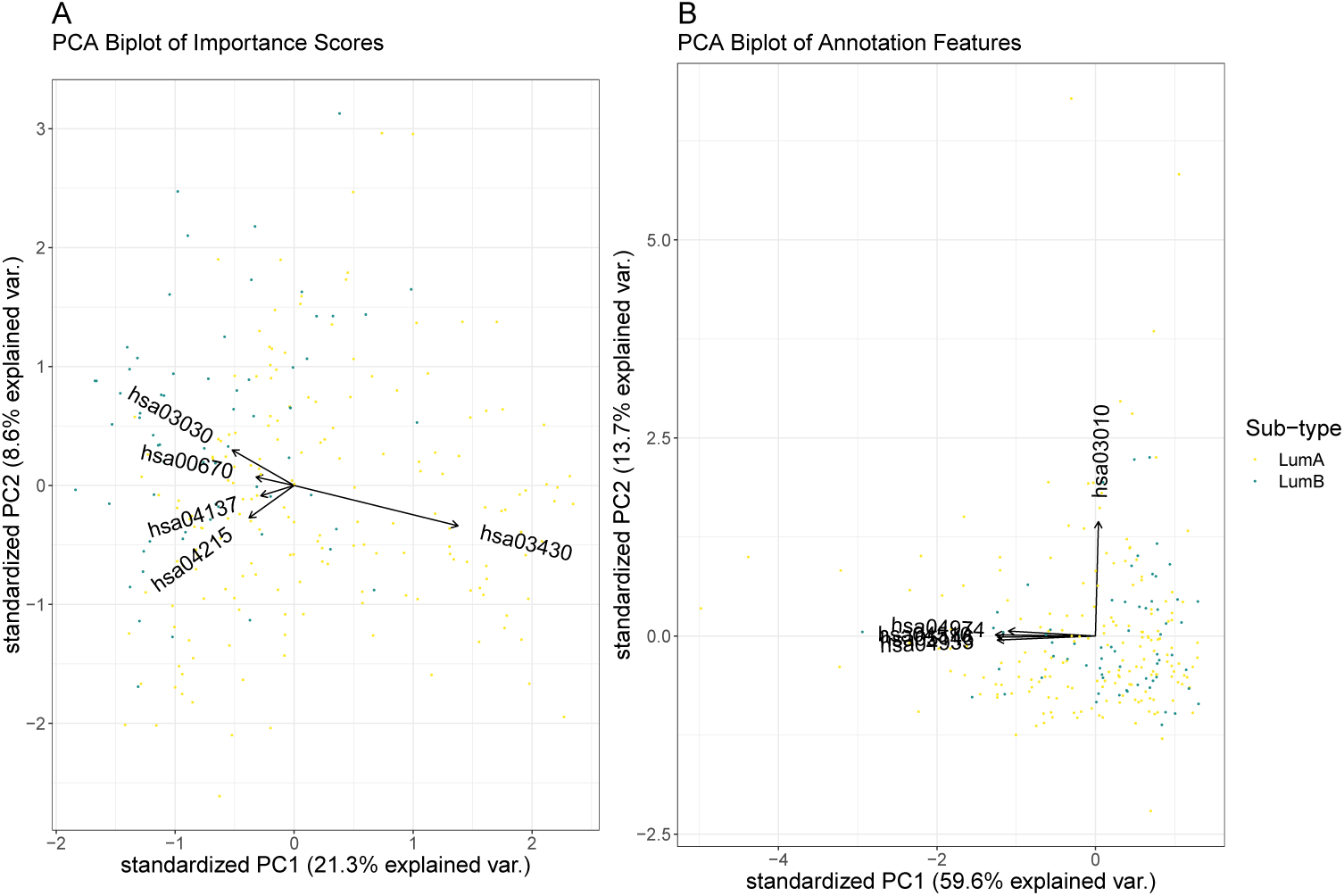
This figure shows a PCA biplot of the KEGG-based importance scores (left panel) and the KEGG annotation features (right panel), with the top 5 most variable terms labelled explicitly. For the importance scores, we see that the first principal axis describes much of the variance between the breast cancer sub-types, while the second principal axis describes much of the variance within the luminal A sub-type. By super-imposing the features as arrows, we can see which annotations best describe the origin of this variance. This level of structure is not evident when looking at the PCA biplot of the annotation feature space. This figure is produced using the test set only.

## 4 Summary

Breast cancer is a complex heterogeneous disorder with many distinct molecular sub-types. The luminal breast cancer class, comprised of the luminal A and luminal B intrinsic sub-types, varies in disease severity, prognosis and treatment response (Dai *et al*. (2015)), and has been described as existing along a vast phenotypic continuum of severity (Netanely *et al*. (2016)). Stratifying individual cancerous samples along this severity continuum could inform clinical decision-making and generate new research hypotheses. In this manuscript, we propose the `DeepTRIAGE` architecture as a general solution to the classification and stratification of biological samples using gene expression data. To the best of our knowledge, this work showcases the first application of the attention mechanism to the classification of high-dimensional gene expression data. In developing DeepTRIAGE, we also innovate the attention mechanism so that it extends to high-dimensional data where there are many more features than samples. Using DeepTRIAGE, we show that the attention mechanism can not only classify cancer sub-types with good accuracy, but can also provide individualised biomarker scores that reveal *and describe* the heterogeneity within and between cancer sub-types. By applying `DeepTRIAGE` to the gene expression signatures of luminal breast cancer samples, we identify canonical cancer pathways that differentiate between the cancer sub-types and explain the variation within them, and find that some of this intra-class variation associates with tumour severity.

